# Evaluating sickness-induced anxiety versus lethargy at the behavioral and neuronal activity level

**DOI:** 10.1101/2022.10.03.510687

**Authors:** Hunter T. Lanovoi, Rumi Oyama, Ioana Carcea

## Abstract

In mammals, inflammatory responses to infections trigger adaptive behavioral changes collectively known as ‘sickness behavior’. Among these, lethargy protects the sick individual by conserving energy, and increased anxiety is believed to prevent exposure to threats. However, the characterization of these conflicting behavioral states in sickness could be an artifact of behavioral assessment, particularly in rodents. We adjusted existing behavioral testing and designed a new paradigm to disambiguate between increased lethargy versus increased anxiety. Our data indicate that in mice sickness induces a significant increase in lethargy but not in anxiety. Further supporting our behavioral results, at the neuronal level we found evidence that sickness activates anxiolytic rather than anxiogenic regions of the amygdala, including oxytocin receptor expressing neurons. Putative mechanisms by which sickness could activate CeA-OTR+ neurons were investigated.

## Introduction

Mammals fight infections by mounting an inflammatory immune response, which can often lead to temporary impairments in physical and cognitive abilities (Dantzer and Kelly 2007; Biesmans et al. 2013; Habba et al. 2012). Sick individuals change their behavior in order to conserve energy for fighting the infection, and to protect themselves from exposure to additional threats. The collection of such protective behavioral changes is known as ‘sickness behavior’, and can be described as a general decrease in activity, or lethargy (Ilanges et al., 2022). Several studies reported that sickness also increases anxiety in rodents (Lacosta et al., 1999; Swiergiel and Dunn, 2007; Kinoshita et al, 2009), a state of high alert and restlessness that is energetically taxing, and differs substantially from lethargy. However, these findings could be confounded by the fact that rodent anxiety tests depend on measures of mobility. In this study, we designed behavioral approaches to disentangle anxiety from lethargy during sickness, and to determine whether sickness is indeed anxiogenic.

In addressing the anxiety phenotype during sickness, we also investigated neuronal activity in brain structures known to play important roles in anxiety regulation. The central amygdala (CeA) has been shown to play essential roles in controlling anxiety responses in humans, non-human primates and rodent models (Oler et al. 2016; Fox et al. 2015; Tye et al. 2011). Increased activity within CeA associates with higher anxiety, and lesions of this structure lead to anxiolytic effects and higher risk-taking behavior (Adolphs et al. 1992; Kalin et al. 2004; Yarkoni et al. 2011). The CeA is composed of 3 subnuclei: medial (CeM), lateral (CeL), and capsular (CeC). Activity of CeM neurons increases fear and anxiety (Ciocchi et al. 2010). On the contrary, activity of neurons in CeL and CeC, most of which are inhibitory neurons projecting to CeM, decreases anxiety (Tye et al, 2011). In particular, oxytocin neuromodulation in CeC and CeL has a marked anxiolytic effect (Viviani et al. 2011; Knobloch et al. 2012). The oxytocin receptor (OTR) is expressed in a subpopulation of CeL and CeC neurons (Haubensak et al. 2010). OTR agonists increase neuronal activity in CeL and CeC inhibitory neurons, and subsequently suppress CeM activity (Huber et al., 2005; Viviani et al., 2011). Although previous work showed that LPS-induced sickness can increase activity in CeA (Marvel et al. 2004; Osterhout et al., 2022, Ilanges et al., 2022), it remains unclear which subnuclei respond. Given the opposite roles of CeM and either CeL/CeC on anxiety, knowing the spatial distribution of neuronal activity changes is critical for understanding the anxiety behavioral manifestation of sickness.

## Results

### Sickness increases immobility and decreases exploration

In rodents, systemic administration of isolated bacterial toxins such as lipopolysaccharide (LPS) can reproduce physiological (fever) and behavioral responses during sickness (Osterhaus et al 2022; Ilanges et al., 2022). Using LPS, previous literature in rodents reported increased anxiety during sickness (Lacosta et al., 1999; Swiergiel and Dunn, 2007; Kinoshita et al, 2009) as early as two hours after administration. However, it remains unclear if these changes reflect increased anxiety after LPS or increased lethargy (Ilanges et al., 2022). To address this, in our first experiment we habituated adult male mice to the experimenter and behavioral apparatus for 2 consecutives days prior to testing behavior (**Figure 1A**), in order to dampen novelty-induced anxiety. On the third day, in half of the mice we induced an inflammatory response by systemically administering 0.5 mg/kg of lipopolysaccharide (LPS). This concentration has been previously shown to induce sickness behavior in mice (Haba et al. 2012; Ilanges et al., 2022). Behavioral performance was tested in the open field and on the elevated plus maze two hours following either saline (N = 8 mice) or LPS (N = 8 mice) injections (**Figure 1A,B,D**).

**Figure 1:**
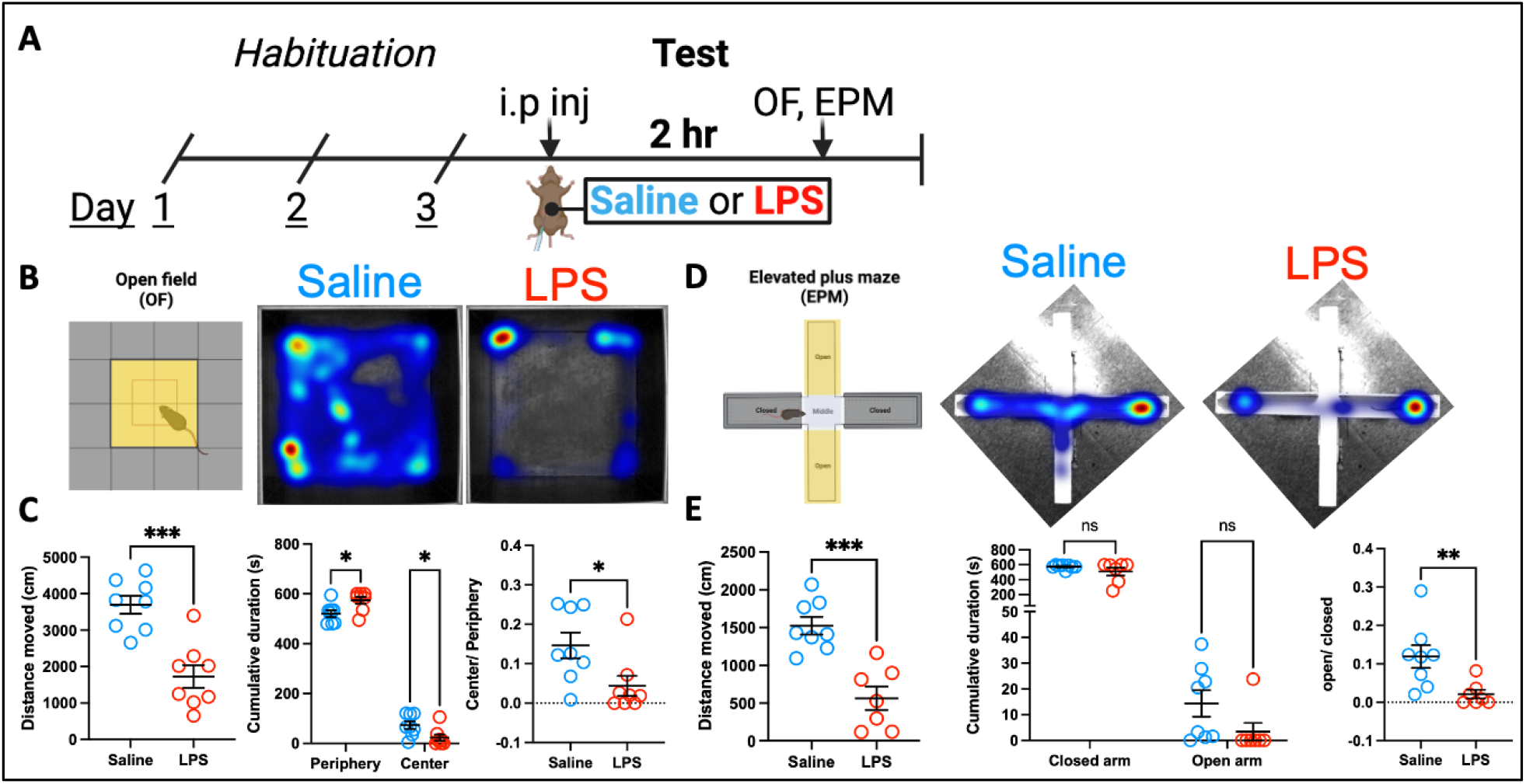
Assessment of open field and plus maze anxiety behavior in sick mice. Paradigm for habituation, LPS or saline treatment, and behavior testing (A). Open field assay design and representative heatmaps (B). Summary data collected from open field assay (C). Elevated plus maze design and representative heatmaps (D). Summary data collected from elevated plus maze (E).

Supporting the lethargic effect of sickness, in the open field task we found that compared to saline treated animals, LPS treatment decreases the total distance travelled (**Figure 1C, left;** ‘Saline’ = 3695 ± 249 cm, ‘LPS’ = 1727 ± 313 cm, unpaired Cohen’s d = −2.46 [95%CI: −3.97, −1.04], unpaired two-tail t-test, p = 0.0002). Sickness also increased thigmotaxis, as LPS treated mice preferred to spend more time in the periphery and less time in the center of the open field compared to saline treated mice (**Figure 1C, center;** ‘Saline periphery’ = 520 ± 13.94 s, ‘LPS periphery’ = 574 ± 13.6 s, unpaired Cohen’s d = 1.38 [95%CI: −0.009, 3.01]; ‘Saline center’ = 73.3 ± 15.15 s, ‘LPS center’ = 22.78 ± 12.65 s, unpaired Cohen’s d = −1.28 [95%CI: −2.7, 0.03]; two-way ANOVA shows an effect of interactions, F(1,14) = 7.225, p = 0.017; Sidak’s multiple comparison test shows significant differences between ‘Saline periphery’ and ‘LPS periphery’, p = 0.021, and between ‘Saline center’ and ‘LPS center’, p = 0.03). Consequently, the ratio of time spent in center compared to time in periphery is smaller in LPS treated mice compared to saline treated mice (**Figure 1C, right;** ‘Saline’ = 0.14 ± 0.03, ‘LPS’ = 0.04 ± 0.02, unpaired Cohen’s d = −1.24 [95%CI: −2.55, 0.04], Mann-Whitney test, p = 0.02).

In the elevated plus maze, we also find that mice move less after LPS injections compared to saline treated animals (**Figure 1D,E left;** ‘Saline’ = 1525.79 ± 117.2 cm, ‘LPS’ = 500.08 ±148.01 cm, unpaired Cohen’s d = −2.74 [95%CI: −3.62, −1.91], unpaired t-test, p < 0.0001). We did not find significant differences between saline and LPS treated animals in terms of time spent in the open and closed arms or the maze (**Figure 1E, middle;** ‘Saline’ closed arms = 574.21 ± 10.12 s, ‘LPS’ closed arms = 457.74 ± 68.93 s, unpaired Cohen’s d = −0.83 [95%CI: −1.72, 0.13], ‘Saline’ open arms = 14.35 ± 5.15 s, ‘LPS’ open arms = 2.98 ± 2.98 s, unpaired Cohen’s d = −0.95 [95%CI: −2.08, 0.18], two-way ANOVA for arms x treatment interaction F(1,14) = 2.18, p = 0.161). When the duration in open arms was normalized to the duration in closed arms, we did find that saline treated animals spent proportionally more time in open arms than LPS animals (**Figure 1E, right;** ‘Saline’ = 0.11 ± 0.02, ‘LPS’ = 0.01 ± 0.01, unpaired Cohen’s d = −1.6 [95%CI: −2.57, −0.58], Mann-Whitney test, p = 0.005).

Given that animals had been habituated to the testing conditions, the observed changes in behavior after LPS treatment are unlikely to result from novelty-induced anxiety. Rather, they can be explained by lethargy (decreased mobility and exploration) during sickness.

To further investigate the link between LPS treatment and anxiety, we tested a separate cohort of twelve mice on a threat exposure experiment. Briefly, after 2 consecutive days of habituation, we tested mouse behavior in response to an intruding robot spider (**Figure 2A**). This threatening stimulus has been previously characterized in group escape behavior in rodents (Kim et al., 2020). We assessed the autonomic response to the robot spider in (n=3 mice) and found a marked increase in heart rate upon introduction of the spider, confirming the perception of threat (**Supplementary Figure 1A**; ‘Baseline’ = 555.36 ± 26.19 beats/min, ‘Spider’ = 686.43 ± 31.82 beats/min, paired Cohen’s d = 2.6 [95%CI: 2.2, 3.17], paired two-sided t-test, p = 0.03). The movement of the robot was remotely controlled by the experimenter, such that it repeatedly approached the mouse (1 approach/min, for 5 minutes). For each approach, we measured the minimum distance from threat that each mouse would tolerate before escaping the spider. Increased anxiety, being associated with risk aversion, would lead to larger distances from threat before escape. Mean distance before escape was not different between LPS and saline treated mice (**Figure 2A,B**; ‘Saline’ = 23.4 ± 1.43 cm, ‘LPS’ = 22.36 ± 3.08 cm, unpaired Cohen’s d = −0.18 [95%CI: −1.47, 1.31], unpaired t- test, p = 0.751). This indicates that the anxiety induced by a potential threat is not different between control and sick animals.

**Figure 2:**
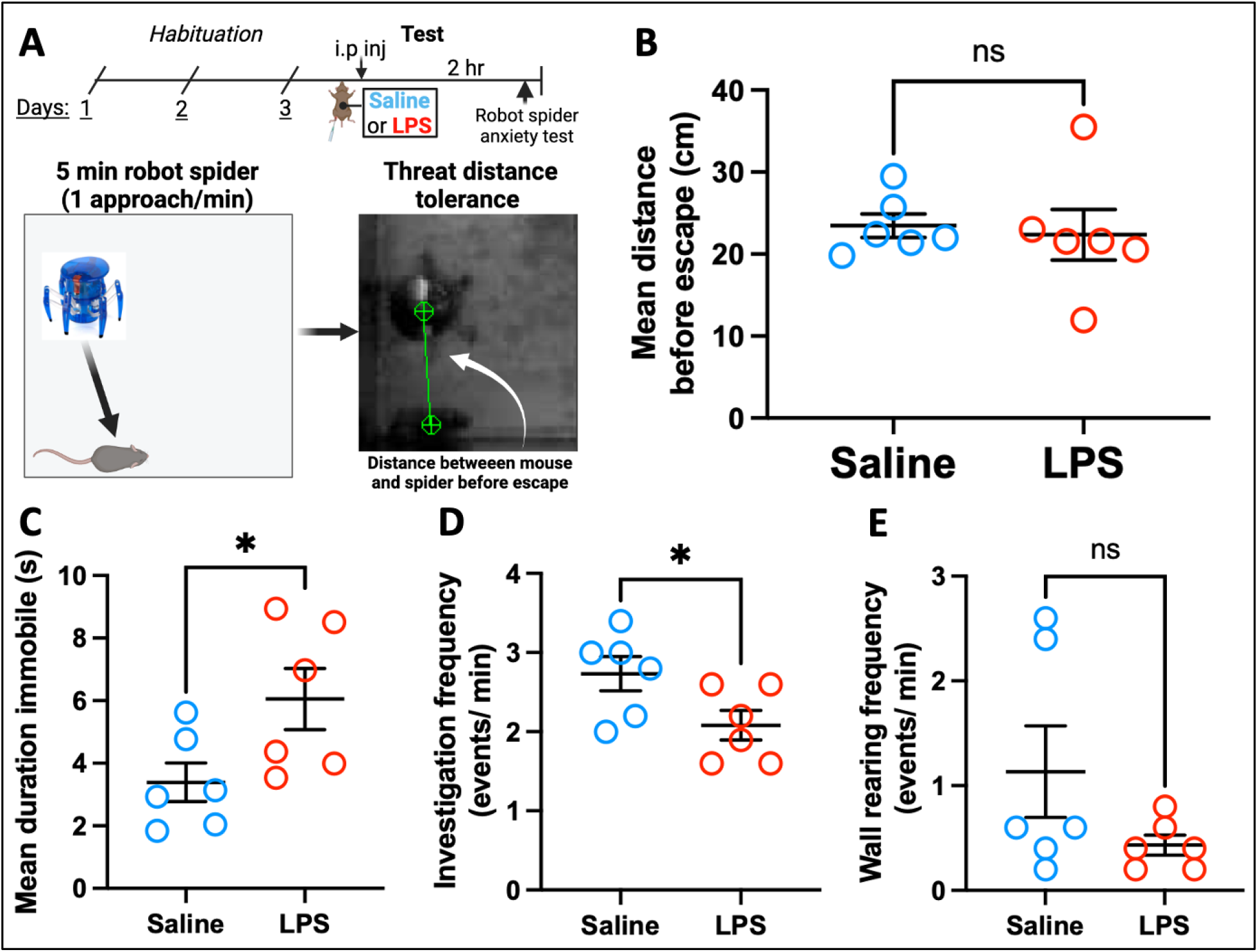
Assessment of anxiety behavior in response to a robot spider threat in sick mice. Paradigm for habituation, LPS or saline treatment, and robot spider approach with measurement of minimum distance before escape (A). Quantification of mean distance in (s) before mouse escape behavior is observed (B). Summary data of experimenter scored behaviors during the 5-minute robot spider interaction (C-E).

Furthermore, additional data from this experiment suggest that LPS increases immobility and decreased exploration: LPS-treated mice spent more time being immobile (**Figure 2C;** ‘Saline’ = 3.38 ± 0.61 s, ‘LPS’ = 6.05 ± 0.97 s, unpaired Cohen’s d = 1.33 [95%CI: 0.15, 2.67], unpaired t-test, p = 0.043) and investigated the spider robot less (**Figure 2D;** ‘Saline’ = 2.73 ± 0.21, ‘LPS’ = 2.08 ± 0.18, unpaired Cohen’s d = −1.31 [95%CI: −2.75, 0.0], unpaired t-test, p = 0.046). Rearing, a behavior that can indicate either anxiety or exploration, was not changed by LPS treatment (**Figure 2E**; ‘Saline’ = 1.13 ± 0.43, ‘LPS’ = 0.43 ± 0.09, unpaired Cohen’s d = −1.31 [95%CI: −2.75, 0.0], Mann-Whitney test, p = 0.29).

### Sickness activates OTR+ neurons in central amygdala

Specific subnuclei and neuronal populations in CeA serve distinct functions in anxiety. Activity of CeM projection neurons increases anxiety manifestations. On the contrary, activity in CeL/CeC, and in particular of OTR+ neurons, has anxiolytic effects. To determine if sickness increases activity of CeA-OTR+ neurons in CeL/CeC, we used six OTR-T2A-TdTomato mice that express a red fluorescent protein in OTR+ cells. After several days of habituation to handling and experimental context, three mice were injected intraperitoneally with saline and three mice with 0.5 mg/kg LPS (**Figure 3A**). Two hours after injection, animals were perfused and subsequently processed for *c-fos* immunostaining. We found that compared to saline, LPS treatment dramatically increases the proportion of active CeA-OTR+ neurons (**Figure 3B,C**; ‘Saline’ = 0.41 ± 0.2%, ‘LPS’ = 16.64 ± 2.6%, unpaired Cohen’s d = 5.07 [95%CI: 3.83, 12.7], unpaired two-tailed t- test, p = 0.003). Importantly, CeA-OTR+ cells represent a substantial proportion of all CeA cells activated by LPS treatment (**Figure 3C, bottom;** ‘Saline’ = 13.42 ± 2.16%; ‘LPS’ = 37.95 ± 3.68, unpaired Cohen’s d = 4.69 [95%CI: 4.14, 5.01], unpaired two-tail t- test, p = 0.004). These results identify CeA-OTR+ cells as major targets for LPS-induced sickness. Furthermore, we find that CeL and CeC host the majority of CeA c-fos+ neurons in LPS treated animals (**Figure 3D;** ‘CeL’ = 111.3 ± 19.33 cells or 33.9%, ‘CeC’ = 242 ± 126 cells or 50.6%, ‘CeM’ = 70.67 ± 50.68 cells or 15.46%). Taken together, our activity studies indicate that LPS increases neuronal activity in anxiolytic CeA subnuclei more than in anxiogenic subnuclei.

**Figure 3:**
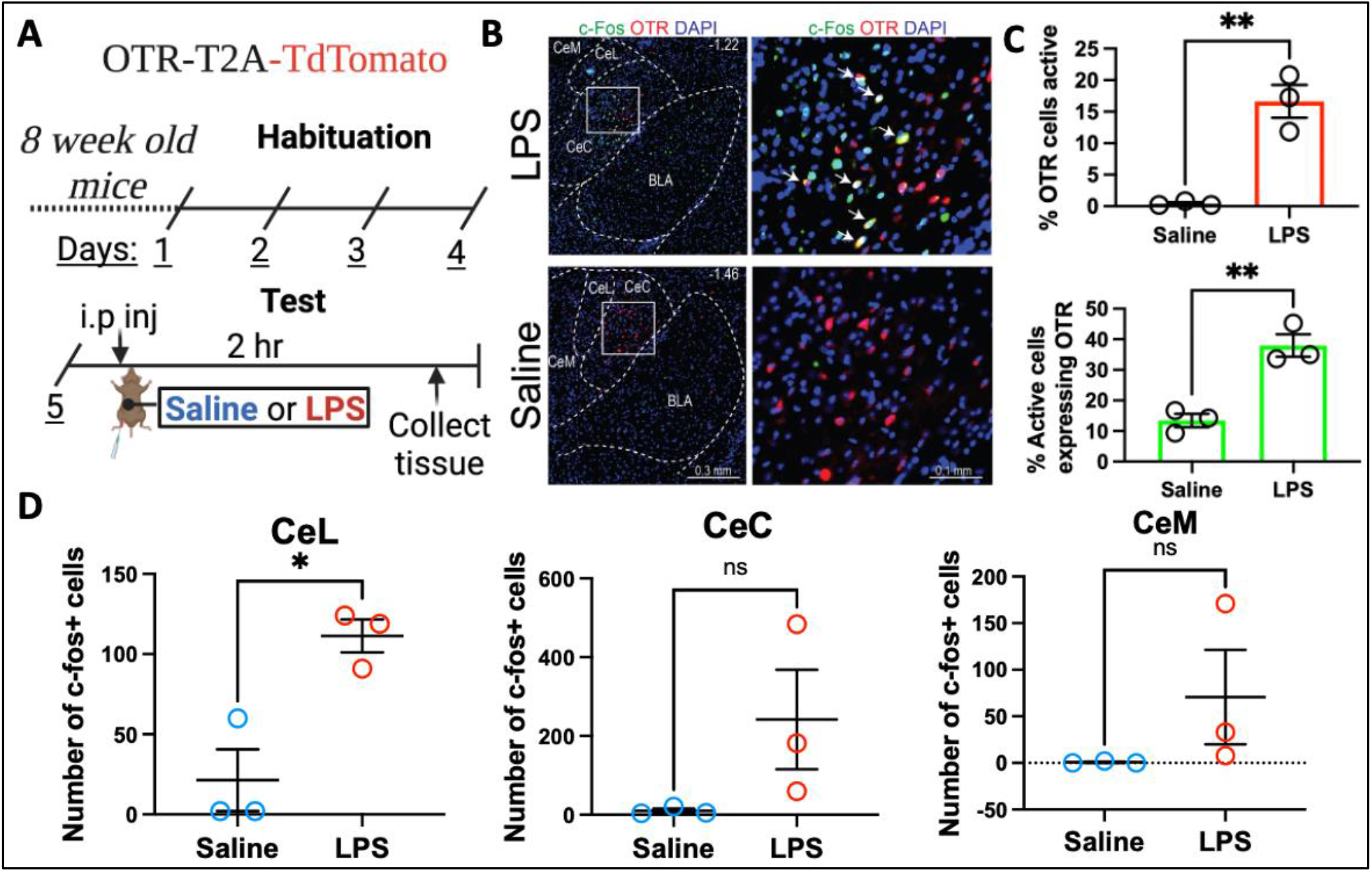
Neuronal activity in CeA of sick OTR-T2A-TdTomato mice. Paradigm for habituation, LPS or saline treatment, and collection of tissue for histological analysis (A). Representative coronal sections in the amygdaloid region of LPS (top) and saline mice (bottom) immunostained for TdTomato and c-fos, and DAPI staining (B). CeA-OTR+ quantification collected from LPS n=3 and saline n=3 mice (C). Distribution of c-fos+ cells in the three subnuclei of the CeA (CeL, CeC, CeM) (D). <u>Abbreviations</u>: CeL: central lateral amygdala CeC: central capsular amygdala CeM: central medial amygdala BLA: basolateral amygdala.

### Potential neuroinflammatory mechanisms by which sickness could increase CeA neuron activity

Sickness can change behavior via the action of inflammatory cytokines on peripheral or central neuronal targets, or as a response to fever. To distinguish between these two possibilities, we monitored core body temperature after LPS and saline injections (Saline n=2, LPS n-5). We confirmed that LPS changes body temperature, with an initial hyperthermia followed by gradual hypothermia (**Supplementary Figure 1B**). At 2 hours after injection, when both the behavioral and neuronal activity studies were performed, core body temperature was similar to pre-injection and to temperature after saline injection. Therefore, it is unlikely that fever could cause the observed changes in behavior. To determine if local neuroinflammation could play a role in inducing c-fos expression in CeA, we performed immunohistochemistry for the transcription factor STAT3. In response to cytokine action, STAT3 translocates to the nucleus where it affects gene transcription and subsequently impacts neuronal excitability (Murase et al. 2012; Han et al. 2021). We detect substantial nuclear localization of STAT3 in CeA following LPS but not saline injections, particularly in CeL and CeC subnuclei (**Figure 4**). This indicates that a potential local neuroinflammatory mechanism could induce increased c-fos expression in the anxiolytic CeA regions.

**Figure 4:**
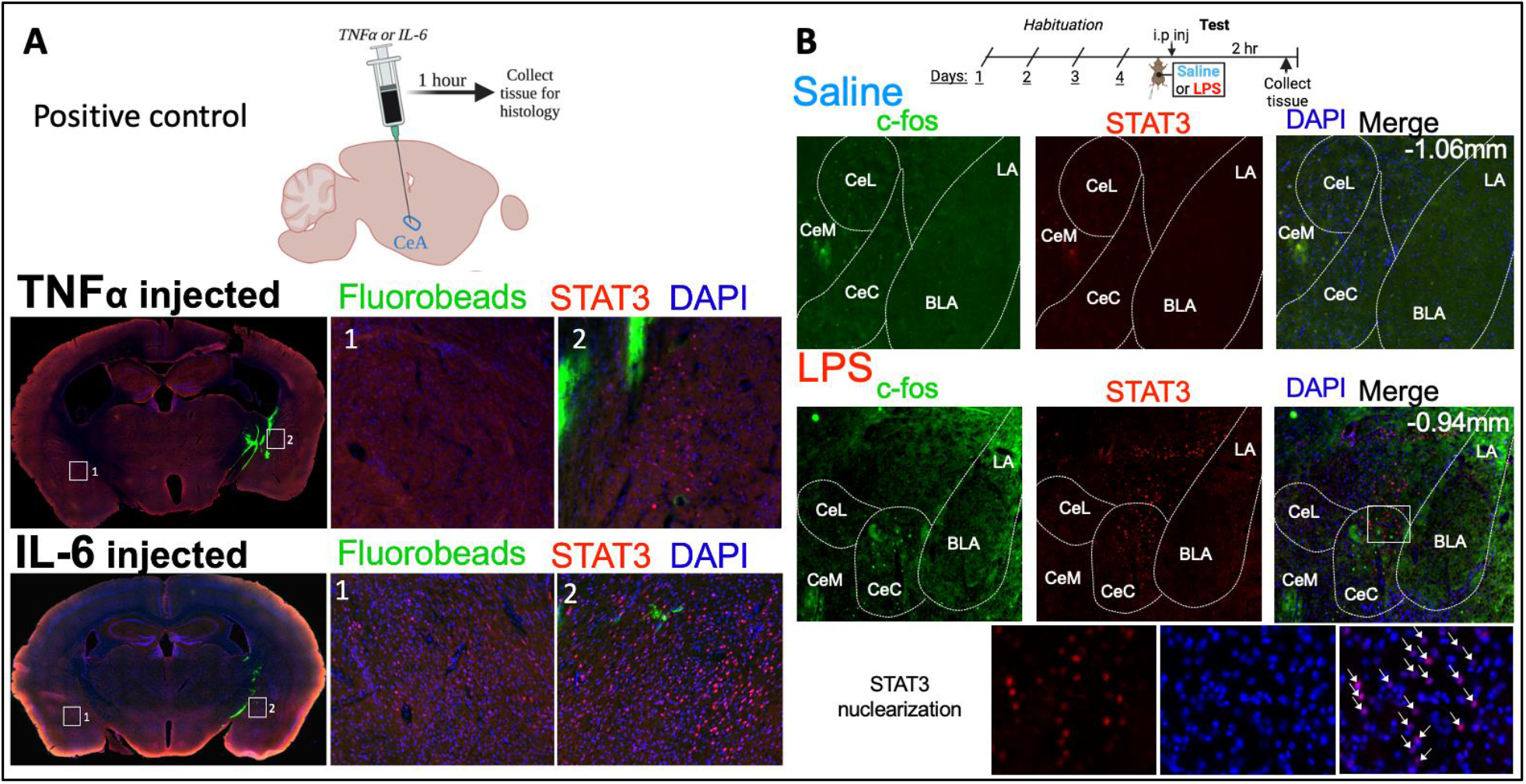
Detection of local neuroinflammation in response to systemic LPS treatment. Paradigm for used to detect cytokine (TNFα or IL6) induced sterile neuroinflammation by staining for nuclear STAT3 (A). Paradigm used to habituate, treat, and collect tissue from mice to determine if systemic LPS induces CeA neuroinflammation (B top). Representative coronal section images in the amygdaloid region of LPS or saline mice stained for nuclear STAT3, c-fos, and DAPI (B middle). Zoomed in image of the CeC with STAT3 nuclear localization. <u>Abbreviations</u>: CeL: central lateral amygdala CeC: central capsular amygdala CeM: central medial amygdala BLA: basolateral amygdala LA: lateral amygdala.

### Potential neural circuit mechanisms by which sickness could increase CeA-OTR+ neuron activity

Sickness can induce c-fos expression in a number of structures along the vagal pathway, including the parabrachial nucleus (Ilanges et al., 2022; Marvel et al. 2004; Campos et al. 2018). In addition, sickness can induce c-fos expression in circumventricular organs in the brain, via direct action of peripheral cytokines (Osterhout et al., 2022), and neuronal projections from these organs can propagate activity to connected brain regions, including the paraventricular nucleus of the hypothalamus. We find active cells (c-fos+) in both the parabrachial nucleus and paraventricular hypothalamus of LPS but not saline treated mice (**Figure 5 A,B**). To determine if sickness and LPS treatment could activate CeA- OTR+ neurons via a neural circuit mechanism, in OTR-T2A-iCre mice we used monosynaptic retrograde rabies tracing from CeA-OTR+ cells. Importantly, we found ipsilateral connections in both the parabrachial nucleus in the brainstem vagal pathway, and in the hypothalamic paraventricular nucleus (**Figure 5**). Many of the neurons in the paraventricular nucleus that projected on CeA-OTR+ cells expressed the oxytocin peptide.

**Figure 5:**
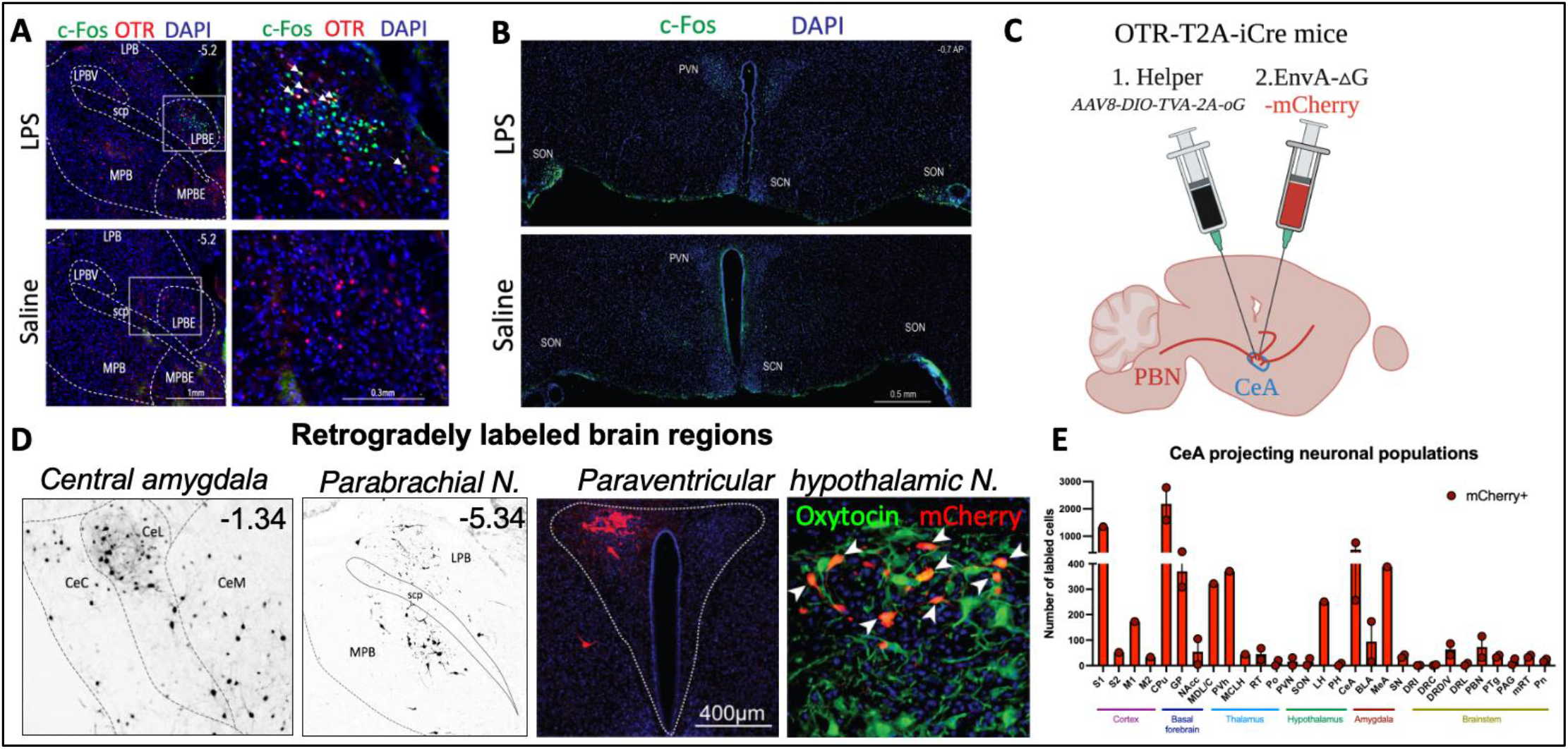
Monosynaptic rabies virus tracing from CeA-OTR+ neurons. Example coronal section around the amygdaloid (A) and paraventricular hypothalamic (B) regions in LPS or saline mice stained for DAPI, c-fos, and tdtomato to mark OTR+ cells. Viral tracing paradigm in OTR-T2A-iCre mice (C). Example coronal images of proximal (CeA) and distal (PBN) retrogradely labeled cells (D left). Representative coronal section images of a population of retrogradely labeled cells in the paraventricular hypothalamic nucleus stained for oxytocin (D right). Summary quantifications of retrogradely labeled cells in all brain areas (E). <u>Abbreviations</u>: PBN: parabrachial nucleus LPB: Lateral PBN LPBV: Lateral ventral PBN scp: superior cerebellar peduncle LPBE: lateral external PBN MPB: medial PBN MPBE: medial external PBN PVN: paraventricular hypothalamus SCN: suprachiasmatic hypothalamus SON: supraoptic hypothalamus.

## Discussion

Here, we used traditional and novel behavior measures of anxiety to disambiguate anxiety from lethargy during LPS induced sickness in rodents. While our results from the open field and elevated plus maze indicates that LPS mice spent less time in risky zones, as reflected by increased thigmotaxis and proportionally less time spent in open arms, this coincides with a significant decrease in movement and in exploration. By challenging sick mice with an acute threat, we were able to assay anxiety more directly, finding significantly increased immobility and decreased investigation indicative of lethargy. Importantly, we found no difference in the distance to the threat that sick mice would tolerate before escape compared to control mice. Taken together our results suggest that traditional behavioral measures of anxiety in rodents do not distinctly assess locomotor deficits versus anxiety. Further, our findings suggest that lethargy might serve adaptive functions that predominate over anxiety behavior during sickness.

By assessing correlates of anxiety at the neuronal level we build upon our behavioral results to find that sick, but not control mice have robust activation of anxiolytic CeC/CeL OTR+ neurons in the amygdala. Crucially, anxiolytic but not anxiogenic amygdalar subnuclei were found to be generally activated during sickness. We found putative neuroinflammatory and circuit mechanisms that might drive the activation of CeA- OTR+ neurons in response to sickness. It remains to be determined if these mechanisms are distinct or act together to change CeA-OTR+ neuronal activity. It will also be important to determine if CeA OTR+ cells are necessary and/ or sufficient to drive lethargy or other behavioral changes during sickness.

## Methods

### Experimental model and subject details

Subjects used for wild-type behavior and CeA immunological studies were adult (8-9 weeks old) wild type C57/BL6 mice purchased from Taconic Biosciences (Hudson, NY, USA). For monosynaptic tracing, and central amygdala LPS activity analysis adult OTR- T2A-iCre and OTR-T2A-tdTomato mice used, generously provided by the Inoue lab (Inoue et al. 2022). For wild-type behavioral experiments, animals received at 6 or 7 weeks of age from Taconic were randomly co-housed in groups of four. OTR-T2A-iCre and OTR-T2A-tdTomato littermates were co-housed in groups of two, three, or four. All animals were housed in standard cages (L:35.56 x W:16.51 x H:17.78 cm) with a 12/12 light-dark cycle and food and water provided ad libitum.

All experiments conducted were in accordance with protocols approved by the Institutional Animal Care and Use Committee at Rutgers.

### Viruses, cytokines, and antibodies

#### Viruses

For chemogenetic studies pAAV-hSyn-DIO-hM3D(Gi) (Cat no. 44362) and pAAV-hSyn-DIO-hM3D(Gq) (Cat no. 44361) were ordered from Addgene. For viral tracing studies AAV8-DIO-TVA-2A-oG and EnvA G-deleted Rabies-mCherry were ordered from the SALK institute.

#### Cytokines

For CeA immunological studies we used carrier free recombinant rat IL-6 Protein (Cat no. 506-RL-010/CF) and carrier free Recombinant Mouse TNF-alpha (aa 80-235) Protein (410-MT-025/CF) purchased from R&D Systems.

#### Antibodies

To stain OTR+ neurons in tissue from OTR-T2A-tdTomato mice, we used primary polyclonal rabbit anti-tdTomato from Takara (Cat no. 632496) and secondary anti-rabbit goat Cy3 (Cat no. 111-165-045) from Jackson Immuno Research. To detect c-fos we used primary polyclonal guinea pig anti-cfos from Synaptic systems (Cat no. 226 004) and secondary anti-guinea pig goat 488 purchased from Jackson Immuno Research (Cat no. 106-545-003). To detect changes in IL-6 signal transduction we used primary monoclonal rabbit anti-Stat3α (D1A5) from Cell Signaling Technologies (Cat no. 8768S) and used the same secondary anti-rabbit antibodies described above.

### Stereotactic surgeries

For all surgeries sterilized surgical instruments were used. Ocular lubricant was used to moisten the eyes and hair on the scalp was removed with Nair hair removal creme. After being placed into the stereotaxic apparatus (David Kopf Instruments), the scalp was disinfected with Povidone-iodine, incised, and retracted. Mice were anesthetized with isoflurane (6% induction, 1% for maintenance). Two coordinates were used for CeA injections and are as follows #1:(AP: −1.52, ML: 2.6, DV: 4.53), #2:(AP: −1.06, ML: 2.35, DV: 4.56). The mice were kept on a heating pad until they recovered from anesthesia and administered SR-buprenorphine (Ethiqa XR) at a dose of 0.5mg/kg before being returned to their home cages

#### Chemogenetics

OTR-T2A-iCre mice received bilateral intracranial injection of pAAV-hSyn-DIO-hM3D(Gi) or pAAV-hSyn-DIO-hM3D(Gq) (200nl) at both sites (#1 and #2) in the left and right CeA (4 total injections). The animals were allowed to recover in their home cage at least 9 days prior to behavioral testing.

#### Neuroinflammation positive control

Wild type C57/BL6 mice received unilateral injections in the CeA (AP: −1.52, ML: 2.6, DV: 4.53) of recombinant TNF-alpha or IL-6 (400nl each of a 5ng/μl solution with yellow green Flurobeads). After injection animals were kept anesthetized under 1% isoflurane for 1 hour at which time animals were transcardially perfused and brain tissue collected.

#### Viral tracing

OTR-T2A-iCre mice received unilateral intracranial injection of AAV8-DIO-TVA-2A-oG (50 nl) in the right CeA (AP: −1.52, ML: 2.6, DV: 4.53). Two weeks following this injection the same mouse received an injection of EnvA G-deleted Rabies-mCherry (200nl) at the same site in the right CeA. Two weeks after the injection of rabies virus mice were sacrificed, perfused and brain tissue collected for subsequent histological analysis.

### LPS activity study

OTR-T2A-Tdtomato mice were habituated to the experimenter, an i.p. saline injection, and the apparatus used for testing for 4 days prior to the test day. On the test day mice were injected with LPS (0.5mg/kg) or Saline and returned to their home cage for the 2 hours before sacrifice and perfusion to collect brain tissue.

### Anxiety behavior testing

C57/BL6 or OTR-T2A-iCre mice were habituated to the experimenter, an i.p. saline injection, and the apparatus used for testing for 2-3 days prior to the test day. On the test day mice were injected with LPS (0.5mg/kg) or Saline and returned to their home cage for the 2 hours before the elevated plus maze and open field tests, the order of which was counterbalanced across groups. For DREADD injected OTR-T2A-iCre mice animals received an injection of deschloroclozapine (DCZ: Sigma) at a dose of 100μg/kg or vehicle 30 minutes prior to behavior testing. Both behavioral tasks were 10 minutes in length and were conducted by a blinded experimenter.

#### Elevated plus maze

The arena used for testing was 55.88 centimeters in length per arm with 12.7-cm arm width at a height of 87.63 centimeters from the ground.

#### Open field

The arena used for testing was L/W/H: 62.23 cm in size and was painted black.

#### Robot spider anxiety test

C57/BL6 mice were habituated to the testing conditions and apparatus the same for open field testing. On the test day mice were injected with LPS (0.5mg/kg) or Saline and returned to their home cage for the 2 hours before testing. The behavioral task was 15 minutes in length and consisted of a 5-minute period before and after a 5-minute robot spider interaction (HEXBUG spider: 451-1652). For the robot spider interaction, a square partition (L/W/H: 20.5 cm) was placed in the center of the open field arena with the spider placed in the periphery. During the 5-minute interaction a blinded experimenter remote controlled the spider to approach the mouse twice per minute. After the 5-minute interaction the robot spider and square partition were removed from the open field apparatus.

### Behavioral analysis

To quantify the movement and duration spent in zones within our apparatuses we used Ethovision XT behavioral tracking software from Noldus (Noldus 2022). Raw heatmaps and trial statistics, including the duration spent in the middle, open, closed arms or center and periphery, were collected, and used for analysis. For the robot spider anxiety test data was time binned by 5 minutes and thigmotaxis was compared before and after the robot spider interaction. Videos collected during the 5-minute spider interaction were further analyzed by an experienced and blinded experimenter using Boris software (Boris 2022) to score threat and anxiety related behavior.

### Histology, immunohistochemistry, and microscopy

Mice were anesthetized with an overdose of i.p. Ketamine (100mg/kg)/ xylazine(10mg/kg) and were then transcardially perfused with 1xPBS, followed by fixation with 4% paraformaldehyde (PFA; Sigma) in 1xPBS. Brain tissue was then carefully extracted and post-fixed in 4% PFA for 2 hours before cryoprotection in 15% sucrose and 30% sucrose each overnight. Coronal slices (25 μm) were collected using a cryostat microtome (Lecia, Germany), and mounted on charged slides using 1xPBS.

#### Staining

Mounted sections were incubated with 1xPBS, containing blocking solution (0.001% TritonX-100 and 5% fetal bovine serum, FBS in 1xPBS, 2 h at room temperature). Primary antibodies (rabbit anti-tdTomato, 1:500; guinea pig anti-cfos, 1:500; rabbit anti-Stat3α, 1:500) were diluted in 1xPBS and applied to tissue samples for a 48-hour incubation at 4°C. The secondary antibodies (anti-guinea pig goat 488, 1:500; anti-rabbit goat 488, 1:500; anti-rabbit goat Cy3, 1:500) were diluted in 1xPBS and used to incubate sections at room temperature for 2 hours. Sections were then incubated with DAPI for 5 minutes before a wash in 1xPBS and cover slipping with Flouromount-GT. All images were photographed and analyzed with a Nikon Ti2 epifluorescence microscope and further processed using ImageJ software.

### Quantification and statistical analysis

Descriptive and inferential statistics were performed in GraphPad Prism (GraphPad Software, Inc.). Estimation statistics (Cohe’s d effect size) were computed as recommended (Ho et al., 2019) with the online platform https://www.estimationstats.com.

## Author contributions

HL and IC designed experiments. HL performed all experiments except the telemetric recordings which were performed by RO. IC and HL wrote the manuscript.

## Acknowledgements

We would like to thank Dr. Steve Levison, Dr. Martin Blaser, Dr. Vanessa Routh for their insightful comments on our experiments. We thank Dr. Annie Beuve for access to the radiotelemetric system for recording core body temperature and heart rate. We thank the Rutgers NJMS vivarium and microscopy core for their support.

---

**Supplemental figure 1:**
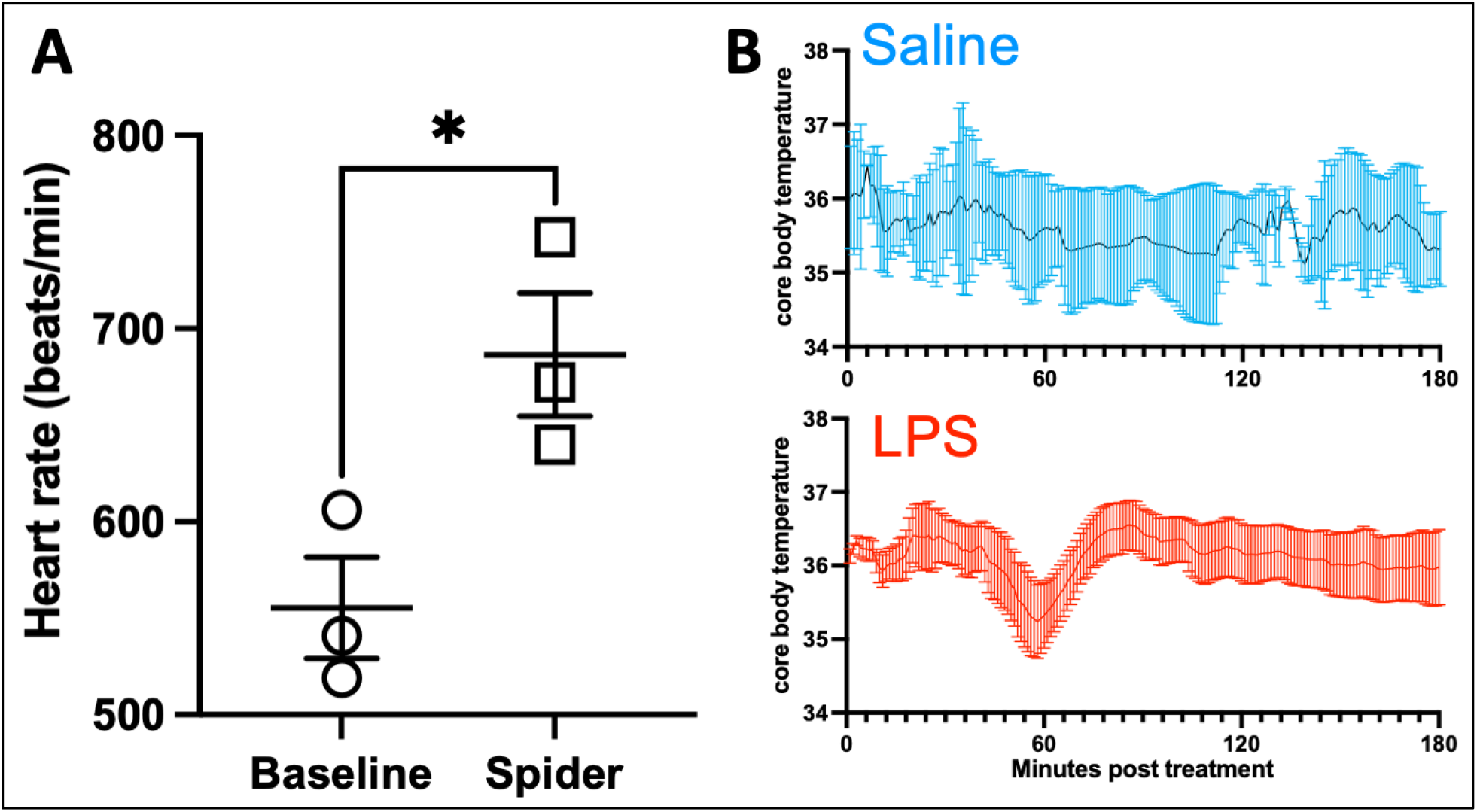
Radiotelemetric recordings of heart rate (beats/min) at baseline (30 min) and after introduction of a robot spider threat (15 min) in n=3 mice (A). Radiotelemetric recordings of body temperature in mice for three hours after treatment with saline or LPS.

